# Environmental history shapes host-associated dynamics of sporulating and non-sporulating bacterial subpopulations during infection

**DOI:** 10.64898/2026.07.21.739833

**Authors:** Hasna Toukabri, Christophe Buisson, Yves Le Vern, Alix Sausset, Mickaël Bourge, Leyla Slamti

## Abstract

Host-associated environments represent ecological contexts that can structure microbial population dynamics, yet their effects on sporulating pathogens remain poorly understood. We investigated how passage through a natural insect host shapes population-level traits in the entomopathogen *Bacillus thuringiensis*. Using *Galleria mellonella* larvae, we compared the characteristics of bacterial populations extracted from insect cadavers with those maintained under *in vitro* conditions. Passage through the host generated a distinct population structure, characterized by the stable coexistence of sporulating and non-sporulating bacteria and a larger non-sporulating fraction than in *in vitro* cultures. Host-extracted bacteria exhibited a different morphology and higher virulence than *in vitro*-grown populations, the latter being largely due to the non-sporulating fraction of the population, as shown by reinfection experiments with each subpopulation isolated *via* fluorescence-activated cell sorting. On the other hand, all subpopulations persisted similarly in the host and completed the infection cycle. Host-extracted subpopulations also showed increased tolerance to oxidative stress, consistent with an adaptation to conditions encountered within insect cadavers. Furthermore, competition assays revealed that non-sporulating bacteria from insect cadavers outcompeted sporulating cells, whereas the opposite was observed for *in vitro*-grown bacteria. In addition, spores produced in the host displayed reduced heat resistance but germinated more efficiently than laboratory-derived spores, highlighting environment-dependent properties which may affect transmission potential. Together, these results demonstrate that the host-associated ecological context drives functional differentiation within bacterial populations and modulates key traits linked to survival, competition, stress tolerance and persistence, emphasizing the importance of host-associated environments in structuring ecological properties of sporulating pathogens.

## Introduction

Bacterial populations are constantly exposed to fluctuating environments that challenge their survival. One strategy to cope with such variability is the emergence of phenotypic heterogeneity within genetically identical populations. This heterogeneity allows subpopulations to adopt distinct physiological states, increasing the probability that at least a fraction of cells survives unpredictable stresses. Functional variability has been shown to enhance survival under diverse conditions, including antimicrobial stress, oxidative stress, and nutrient limitation [1, 2, 3]. Environmental history, stress intensity, and local interactions between cells shape the proportion of stress-prepared individuals and influence growth resumption dynamics, highlighting phenotypic diversification as a key adaptive strategy balancing growth and persistence [4, 5].

In sporulating bacteria, the formation of endospores represents an exceptional survival strategy, allowing cells to withstand harsh environmental conditions. This process is tightly regulated, with Spo0A acting as a master regulator that governs entry into sporulation and drives the differentiation of cells into distinct developmental fates in the model bacterium *Bacillus subtilis* in laboratory conditions [6, 7, 8]. Activation of Spo0A is heterogeneous, generating only a fraction of cells that commit to sporulation even under favorable conditions [9]. Positive feedback within the phosphorelay supports intergenerational memory and a bet-hedging strategy during nutrient limitation [10]. As a result, sporulating and non-sporulating cells coexist, providing flexibility in adaptation to fluctuating environments.

Importantly, survival is not restricted to sporulation. Non-sporulating *Bacillus* cells persist under adverse conditions and may gain selective advantages. Asporogenic *Bacillus cereus* variants with *spo0A* mutations survive in oligotrophic environments and display altered metabolic strategies promoting persistence [11]. Reduced sporulation in *Bacillus thuringiensis* has also been associated with diversification and adaptation in multispecies communities [12] and oligosporogenic mutants can retain or enhance insecticidal activity [13]. In addition, non-sporulating *B. subtilis* can survive deep starvation in a slow-growing oligotrophic state [14]. Moreover, vegetative cells impaired in sporulation can persist in food environments while remaining pathogenic [15]. Together, these findings highlight the adaptive value of coexistence between sporulating and non-sporulating phenotypes.

The entomopathogenic bacterium *B. thuringiensis*, a member of the *B. cereus* group, exhibits a complex ecology and remarkable adaptability. This group includes both environmental bacteria and pathogens with versatile lifestyles [16]. *B. thuringiensis* is found ubiquitously in the soil and insects are a major niche for its replication and transmission [17, 18]. Its ecology is dynamic, shaped by interactions with diverse hosts and environmental contexts [19] including amoebae or plant roots, illustrating its capacity to adapt through diverse developmental pathways [20, 21]. Within insect hosts, *B. thuringiensis* infection cycle is highly context-dependent, with germination, growth, and sporulation occurring only under permissive host conditions [22, 23]. During infection, *B. thuringiensis* complete a full infection cycle controlled by quorum-sensing systems involving virulence, necrotrophism, and sporulation [24]. Single-cell analyses revealed heterogeneity within the population during infection with the coexistence of sporulating, necrotrophic, and undifferentiated cells [25]. Additional studies showed that bacterial populations recovered from insect cadavers contain living non-sporulating cells [25]. These cells persist for an extended period of time, present a slowed-down metabolism and exhibit an enhanced oxidative stress response [26]. However, the impact of environmental history on the physiological properties of sporulating and non-sporulating subpopulations remains poorly understood. In particular, it is unclear how passage through a host influences the differenciation and physiological properties of the bacteria.

We therefore investigated how a 7-day passage through *G. mellonella*, a natural host of *B. thuringiensis* particularly suited for studying bacterial physiology, shapes subpopulations dynamics. Through total population and FACS-sorted subpopulations assays using a fluorescent sporulation reporter, we monitored sporulating (Spo^+^) and non-sporulating cells (Spo^-^) using the red fluorescent sporulation reporter P*spoIIQ’mcherry* . These subpopulations were compared depending on whether they originated from insect cadavers at 7 days post-infection (7 dpi) or laboratory-grown cultures in LB during 7 days (7 dLB). Spo^-^ cells extracted from insect cadavers were shorter and more virulent than LB-grown cells. Competition assays showed that Spo^-^ 7 dpi cells outcompete Spo^+^ 7 dpi cells, whereas the opposite occurs in laboratory-grown subpopulations. However, *in vitro*- and insect-extracted subpopulations persisted similarly and completed the canonical infection cycle with partial sporulation. In addition, 7 dpi cells displayed enhanced tolerance to H₂O₂, and 7 dpi spores germinated faster than 7 dLB spores and showed lower heat resistance.

## Material and methods

### Bacterial strains and growth conditions

The acrystalliferous *Bacillus thuringiensis* 407 Cry⁻ strain [27] was used as the parental strain of all strains used in this study. *Escherichia coli* DH5α [28] was used for plasmid construction, and *E. coli* ET12567 [29] was used to prepare DNA prior to electroporation into *B. thuringiensis*. Cells were grown with shaking at 30°C or 37°C in LB medium (1% tryptone, 0.5% yeast extract, 1% NaCl) or HCT medium (0.7% casein hydrolysate, 0.5% tryptone, 0.68% KH₂PO₄, 0.012% MgSO₄·7H₂O, 0.00022% MnSO₄·4H₂O, 0.0014% ZnSO₄·7H₂O, 0.008% ferric ammonium citrate, 0.018% CaCl₂·4H₂O, pH 7.2) [30]. Bacteria were stored at −70°C in LB supplemented with 15% glycerol. When required, antibiotics were added at 100 μg/mL ampicillin for *E. coli* and 10 μg/mL erythromycin for *B. thuringiensis*.

### Plasmid and strain construction

DNA manipulations are detailed in Text S1. All the plasmids, strains, and oligonucleotide primers used in this study are listed in Tables S1a to c.

### Infection of Galleria mellonella

Intrahemocoelic injections were performed using third-instar *Galleria mellonella* larvae (60 ± 10 mg). Equivalent numbers of bacteria (2 × 10³ cells) from three experimental conditions were injected: exponential-phase cells, 7-day post-inoculation in LB cells (7 dLB), or 7-day post-infection insect-extracted cells (7 dpi). For virulence reinfection assays, either the total populations or the sporulating (Spo^+^) and non-sporulating (Spo^-^) subpopulations sorted by FACS were used. Inocula were verified by plating on LB agar. Infected larvae were incubated at 30°C and monitored by time-lapse photography. Larvae were considered dead when movement ceased and melanization occurred.

### Extraction of *B. thuringiensis* from insect cadavers

Insect cadavers were crushed in saline using a FastPrep homogenizer (MP Biomedicals). Briefly, cadavers were placed in 2 mL flat-bottom FastPrep tubes containing five sterile 3 mm glass beads and 1 mL of sterile saline (0.9% NaCl). Samples were homogenized at 6.5 m/s for 60 s. For persistence assays, crude homogenates were used to enumerate the number of total cells by plating on LB agar plates following serial dilutions in saline. For single-cell assays, homogenates were filtered through cotton to remove debris. The filtrate was centrifuged, and the pellet resuspended in saline. When required, cells were fixed in 4% formaldehyde, washed, resuspended in GTE buffer, and stored at 4°C prior to flow cytometric or microscopic analyses.

### Flow cytometric analysis

Bacteria were diluted in filtered saline, and fluorescent events were measured on a CyFlow Space cytometer (Sysmex Partec, France). Specifications of the apparatus and flow cytometric analyses are described in Text S1.

### Microscopy

Bacteria were observed with an AxioObserver.Z1 Zeiss inverted fluorescence microscope with a Zeiss AxioCam 807 mono digital camera and with Zeiss fluorescence filters. mCherry was imaged using the 45 HE filter (excitation: BP 590/20, beam splitter: FT 605, emission: 620/14). Images were processed using the ZEN software package (Zeiss) and ImageJ. Cell length measurements were performed on Spo^-^ populations identified using the P*spoIIQ*’*mcherry* reporter of cells incubated 1 day in LB medium (1 dLB) or extracted from insect cadavers at 1 day post-infection (1 dLB). Individual cells were manually outlined in ImageJ, and the major axis of each cell was measured to obtain cell length. At least 390 cells per condition were measured to calculate average cell lengths and standard deviations.

### Sporulation assay

Fixed bacteria (as described above) harboring the red fluorescent sporulation reporter P*spoIIQ’m-cherry* were analyzed. Red fluorescent events were counted among the total cell count to calculate the percentage of Spo^+^ cells.

### Cell sorting procedure

Spo^+^ cells, Spo^-^ cells and exponential phase cells were sorted with a MoFlo AstriosEQ cell sorter (Beckman-coulter). The instrument used a 70-µm nozzle with a sheath pressure at 60 PSI. Home-made PBS (137 mM NaCl Supelco Ref. 1.06104.1000, 2.7 mM KCl Sigma-Aldrich, Cat# P3911, 8 mM Na2HPO4 Ref. PRO-28028.298, 2 mM KH2PO4 Merck Ref. 1.06579.0500) was used as sheath fluid to avoid bactericide or NaN3 present in commercial sheath liquid. The sorter was calibrated with Ultra Rainbow Calibration particles (Spherotech). Frequency of drop formation was close to 96,000 Hz to reach a flow rate around 40,000 events per second. A first gate was made on a Forward Scatter (FSC-Height) - Side Scatter (SSC-Height) dot plot to select the bacterial population. Doublets were discarded using an SSC-Area – SSC-Height dot plot. For infection cycle analysis and H_2_O_2_ assay, the strain used was Bt (pP*nprA*’*gfp_Bte_AAV*-P*spoIIQ*’*mcherry*) to follow necrotrophism and sporulation upon reinfection (See Table S1). As necrotrophism is no longer expressed in late samples, Spo^-^ cells were characterized as non-fluorescent (Gfp-negative, mCherry-Negative), and Spo^+^ cells as red-fluorescent only (mCherry-positive). For competition assay, the strain used was Bt (P*c*’*gfp_Bte_*-P*spoIIQ*’*mcherry*) which carries a constitutive GFP reporter (P*c*’*gfp_Bte_*, promoter region of the *sarA* gene) in combination with a sporulation reporter, P*spoIIQ*’*mcherry* (See Table S1). The Spo^-^ subpopulation was characterized as green fluorescent cells (GFP-positive, mCherry negative) and the Spo^+^ subpopulation was characterized as red fluorescent cells (mCherry-positive). To collect the fluorescence signal, a 525/52-nm bandpass filter was used for the green fluorescence after excitation with solid blue-laser emitting at 488 nm, and a 620/29-nm bandpass filter for the red fluorescence after excitation with a solid yellow/green-laser emitting at 561 nm. 10^7^ bacteria were collected per condition, samples were then concentrated on 0.22 µm membane filters using a glass filtration apparatus, resuspended in 500 µL saline and plated on LB agar to obtain final bacterial count.

### Hydrogen peroxide tolerance assay

Equivalent numbers of bacteria (approximately 2 × 10³ cells) from three experimental conditions (exponential-phase cultures, 7 dLB cells, and 7 dpi cells) were serially diluted in saline and plated on LB agar supplemented or not with increasing concentrations of H₂O₂ (0.075, 0.150, 0.250, and 0.500 mM). The assay was performed either using total populations or using Spo^+^ and Spo^-^ subpopulations sorted by FACS from the same three conditions. Survival was determined relative to untreated controls.

### Competition assay

To compare the fitness of Spo^+^ and Spo^-^ cells, competition experiments were performed using sub-populations sorted by FACS from 7 dLB cells and 7 dpi cells. Within each condition, Spo^+^ and Spo^-^ bacteria were competed against each other. Spo^-^ cells were isolated from the CS strain, which carries the constitutive GFP reporter P*c*’*gfp_Bte_* in combination with a sporulation reporter P*spoIIQ*’*mcherry* (see Table S1). GFP-positive and mCherry-negative cells were sorted as the Spo^-^ cells. Spo^+^ cells were isolated from the S strain, which carries only the P*spoIIQ*’*mcherry* reporter, and mCherry-positive cells were sorted accordingly. Sorted Spo^-^ cells (GFP⁺) and Spo^+^ (mCherry⁺) cells were mixed at a 1:1 ratio and either incubated in LB for 16 h at 30°C or injected into third-instar *G. mellonella* larvae (2x10^3^ bacteria per larva) and recovered 16 h post-infection at 30°C. The proportions of GFP-positive bacteria (derived from Spo^-^ cells) and non-fluorescent bacteria (NonFluo, derived from Spo^+^ cells that had germinated and lost mCherry fluorescence) were determined. As a control, exponential-phase cells from both reporter strains were sorted and mixed using the same procedure; in this case, GFP-positive cells were competed against non-fluorescent cells, as sporulation had not yet occurred.

The competitive index (CI) was calculated to quantify relative fitness. For 7 dLB and 7 dpi populations, the CI of non-sporulating cells was calculated as:

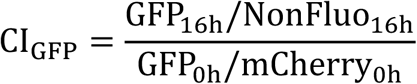

For exponential-phase cells, CI was calculated as:

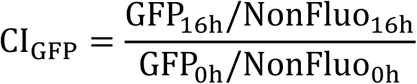

Here, “0h” corresponds to the beginning of the assay (mixing of the bacteria and plating on LB agar). “16h” corresponds to 16 h of co-culture in LB or post-infection recovery from larvae.

### Spore heat resistance assay

Spores were prepared from 7 dLB cultures, 7 dpi cells and 7 day HCT (7 dHCT) cultures similarly to the protocole published in Perchat et al., 2024 [31]. Briefly, the cultures were centrifuged at 5,000 g for 15 min at 4°C to prevent germination and the spore pellet was washed three times with cold distilled water to discard potential remaining germinant, resuspended in cold distilled water then heated for 15 min at 70°C to kill residual vegetative cells. Spore suspensions were then cooled on ice before storage at 4°C until use. For the heat resistance assay, 0.5 mL of the spore suspensions were exposed to 90°C for 0 min, 5 min, 10 min, 30 min and 60 min and quantified by plating onto LB agar. Spore concentrations were expressed as colony-forming units per milliliter (CFU/mL). Survival curves represent log(N/N0) during time, with N0 as initial spore concentration and N as spore concentration after treatment.

### Germination assay

Germination of spores prepared from 7 dLB cultures, 7 dpi cells and 7 dHCT cultures was assessed using the P*spoIIQ*’*mcherry* transcriptional reporter. Spores were incubated in germination buffer (10 mM Tris–HCl, pH 7.4, and 10 mM NaCl) supplemented with L-alanine (10 mM), inosine (1 mM) and tenfold diluted LB medium containing 3% glucose, which allows cell elongation without substantial division. At selected time points (30, 60, and 120 min), the proportion of fluorescent cells (Spo^+^) was quantified by flow cytometry. To account for differences in the initial spore population, the data were normalized to the t = 0 value and proportion of sporulating cells (Spo) was determined as :

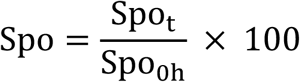

where Spot is the number of mCherry-positive cells at time t, and Spo_0h_ is the number of mCherry-positive cells at the beginning of the assay. This normalization allows comparison of germination kinetics across 7 dHCT, 7 dLB and 7 dpi spores.

### Statistical analysis

The data were analyzed using Prism v8 software (GraphPad).

## Results

### Nonsporulating bacteria are more abundant and display reduced cell length in insect host cadaver compared with *in vitro* conditions

A previous study showed that the bacterial population during infection is heterogeneous and that a fraction of cells can persist in insect cadavers for extended periods in a nonsporulated state [26]. However, whether this heterogeneity is observed across distinct growth environments has not been characterized. We therefore quantified the proportion of sporulating (Spo^+^) cells using strain Bt (pP*spoIIQ*’*mcherry*) and flow cytometry analysis (as described in Material and Methods). Bacteria were recovered from *G. mellonella* insect cadavers at 1, 3 and 7 days post-infection (dpi), and from cultures grown in LB and HCT, a sporulation-inducing medium, at the corresponding time points (Fig. 1a). In insect cadavers, approximately 50% of the bacterial population was mCherry-positive, indicating commitment to sporulation, and this proportion remained stable over time. In contrast, a higher sporulation level was observed under *in vitro* conditions, with approximately 70% and 90% of Spo^+^ cells in LB and HCT, respectively. These proportions also remained stable between 1 and 7 dpi. Together, these results show that the sporulation level differs across conditions, with substantial nonsporulating fractions in both insect cadavers (∼50%) and LB cultures (∼30%).

**Figure 1.**
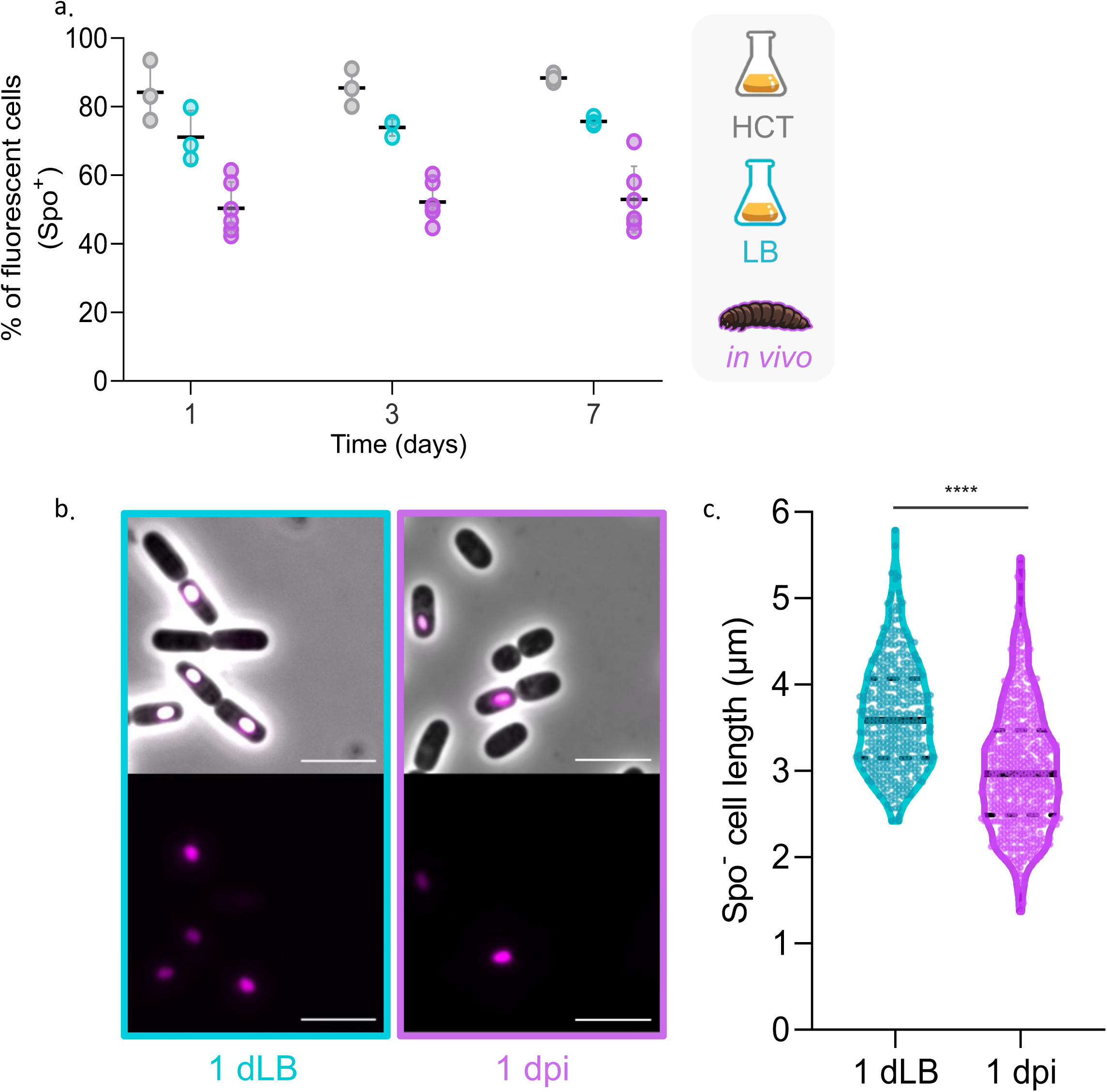
Sporulation quantification and morphological characterization of *Bacillus thuringiensis* populations during insect infection and under *in vitro* growth conditions. (a) Quantification of Spo^+^ cells using strain Bt (pP*spoIIQ*’*mcherry*). Bacteria were recovered from insect cadavers at 1, 3 and 7 days post-infection (purple) or at 1, 3 and 7 days post-inoculation from cultures in LB (blue) or in sporulation-inducing medium HCT (gray) at 30°C. The percentage of fluorescent cells was determined by flow cytometry, as described in Materials and Methods. Each symbol represents the value obtained from one biological replicate. For the insect condition, each symbol corresponds to bacteria recovered from an individual larva (n = 6 larvae, 2 larvae were used per experiment). Three independent experiments were performed. Horizontal black bars indicate the mean, and error bars represent the standard deviation (SD). (b) Microscopy observations of sporulating (Spo^+^) and non-sporulating (Spo^-^) subpopulations. Bacteria carrying the P*spoIIQ*’*mcherry* reporter were observed by epifluorescence and phase-contrast microscopy at 1 day post-inoculation in LB (1 dLB, left pannels) or 1 day post-infection in insect cadavers (1 dpi, right pannels). The top panels show the merge between the phase contrast and epifluorescence images channels; the bottom panels show epifluorescence images. Spo^+^ cells were false-colored in pink based on mCherry fluorescence. Representative images are shown. The scale bars represent 5 μm. (c) Cell length distribution of Spo^-^ bacteria. Cell length (μm) was measured from epifluorescence and phase-contrast microscopy images for Spo^-^ (mCherry-negative) cells recovered at 1 dLB (blue) or 1 dpi (purple). Each dot represents one cell. Violin plots show the distribution of cell lengths, with the median indicated by a black line and quartiles shown as dashed lines. At least 390 cells were measured per condition across three independent experiments. Statistical significance was assessed using a Mann–Whitney test (****, p < 0.0001).

We next assessed the morphology of the nonsporulating subpopulation (Spo^-^). Using the same strain harboring the P*spoIIQ*’*mcherry* reporter to exclude cells engaged in sporulation, cell length was measured using microscopy for Spo^-^ bacteria recovered from insect cadavers at 1 day post-infection and from LB cultures at 1 day post-inoculation (Fig. 1b). Cells isolated from insect cadavers were shorter than cells from LB culture (mean of 3 µm and 3,5 µm, respectively), with an average reduction of approximately 0.5 µm (Fig. 1c). Together, these results show that bacterial populations remain heterogeneous across conditions, with distinct proportions and morphology of nonsporulating cells depending on the environment.

### Prior bacterial environment modulates *Bacillus thuringiensis* virulence during infection without affecting post-mortem persistence or the infection cycle

To investigate whether the observed differences in sporulation distribution and cell morphology were associated with functional changes, we assessed how prior environmental conditions influence bacterial virulence and host persistence. We compared the progression of the infection induced by strain Bt (pP*nprA*’*gfp_Bte_AAV*-P*spoIIQ*’*mcherry)* using exponential phase cells (expo) as a control, 7 days LB culture cells (7dLB), and 7 days post-infection cells (7 dpi) recovered from *G. mellonella* cadavers (Fig. 2). As described in the Material and Methods section, equivalent numbers of bacteria were injected into 3^rd^ instar *G. mellonella* larvae. Figure 2a shows that expo cells killed hosts rapidly, with less than ∼20% survival at 16 hours post-infection (hpi). At 24 hpi, larvae infected with 7 dpi cells showed ∼65% survival, whereas those infected with 7 dLB cells showed ∼75% survival. By 96 hpi, survival dropped below 10% and was about 50% and 70% for larvae infected with expo cells, 7 dpi cells and 7 dLB cells, respectively. These results show that exponential phase cells are more virulent than 7 day-old cells, wether *in vitro*-grown or extracted from insects, and reveal that 7 day-old bacteria recovered from insect cadavers are more virulent than 7 day-old *in vitro*-grown bacteria. We next quantified persistence in insect cadavers of bacteria originating from the different conditions by plating on LB agar (Fig. 2b). Following host death, all bacterial populations, whether the infection was performed with expo, 7 dLB, or 7 dpi cells, persisted at a similar level (∼2.5 × 10^8^ CFU/mL) at 1, 3, and 7 days post-infection.

**Figure 2.**
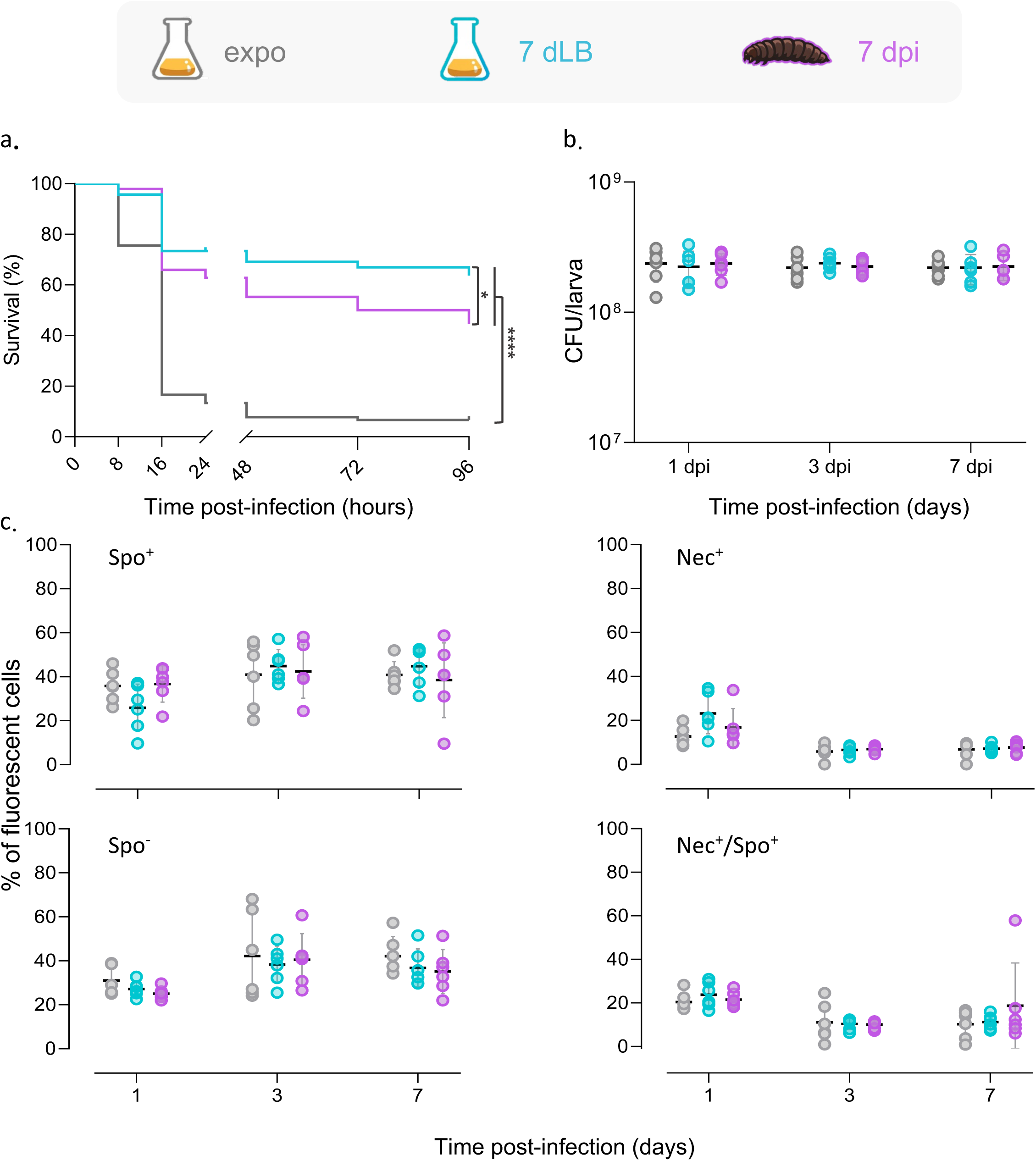
Analysis of virulence, persistence, necrotrophism and sporulation during infection with total bacterial populations. (a) Survival of *G. mellonella* larvae following infection with bacteria derived from *in vitro* and *in vivo* total populations using strain Bt (pP*nprA*’*gfp_Bte_AAV*-P*spoIIQ*’*mcherry*). Exponential-phase cells (gray), cells grown for 7 days in LB (7 dLB, blue), and bacteria recovered from insect cadavers at 7 days post-infection (7 dpi, purple) were injected intrahemocoelically into third-instar larvae (2 × 10³ bacteria per larva). Larvae were incubated at 30°C, and survival was monitored over time (hours post-infection). The data represent three independent experiments with 30 larvae per condition in each experiment. Survival curves were compared using the Gehan–Breslow–Wilcoxon test. Significance is indicated as follows: *, p< 0.05; ****, p < 0.0001. (b) Bacterial persistence in insect cadavers. Following infection with the same populations described in panel a., larvae were crushed at the indicated time points (days post-infection) using a FastPrep homogenizer. Homogenates were serially diluted in saline and plated onto LB agar to determine colony-forming units (CFU/mL). Each symbol represents bacteria recovered from one larva (n = 6 larvae per time point, 2 larvae were used per experiment). Three independent experiments were performed. Horizontal black bars indicate the mean, and error bars represent the standard deviation (SD). (c) Flow cytometry analysis of subpopulations after reinfection with the same populations described in panel a. The strain Bt (pP*nprA*’*gfp_Bte_AAV*-P*spoIIQ*’*mcherry* was used to monitor necrotrophism and sporulation after reinfection. The percentage of fluorescent cells was determined by flow cytometry as described in Material and Methods for Spo^+^ cells (left, top panel), Spo^-^ cells (left, bottom panel), necrotrophic cells (Nec^+^, right, top panel) and Nec^+^/Spo^+^ cells (right, bottom panel) as a function of time (days post-infection) after infection with exponential phase-(gray), 7 dLB (blue), and 7 dpi (purple) populations. Each symbol represents bacteria extracted from one larva (n = 6 larvae per time point, 2 larvae were used per experiment). Three independent experiments were performed. Horizontal black bars indicate the mean, and error bars represent the standard deviation (SD).

To examine whether the bacteria could complete the infection cycle, the same strain was used to track necrotrophism and sporulation (Fig. 2c). At 1 day post-infection, ∼40% of the bacterial population was necrotrophic across all conditions. Then the total number of necrophic bacteria decreased to about 10% of the population and remained at that level until the end of the experiment. Approximately, approximately half of the population entered sporulation, independent of whether the cells originated from expo cells, 7 dLB cells, or 7 dpi cells.

Together, these results show that prior environmental conditions influence bacterial virulence. However, once the host dies, the bacteria from these different environments persist at comparable levels and reproduce the canonical infection cycle, with early necrotrophism and sporulation for part of the population.

### Insect-extracted non-sporulating cells are more virulent than sporulating cells and late LB-cells, with all subpopulations persisting and completing the infection cycle similarly

To assess the contribution of the different bacterial subpopulations to virulence and persistence, using strain Bt (pP*nprA*’*gfp_Bte_AAV*-P*spoIIQ*’*mcherry*), expo, 7 dLB, and 7 dpi cells were sorted by FACS. While Spo^+^ and Spo^-^ subpopulations were sorted out for 7 dLB and 7 dpi populations, expo cells were recovered as a whole population after undergoing the same sorting procedure. Then, equivalent numbers of sorted cells were injected into 3^rd^ instar *G. mellonella* larvae, as described in the Materials and Methods section. As observed with the non-sorted cells experiment (Fig. 2a), expo cells showed rapid host killing, with only ∼10% survival at 24 h post-infection (hpi). For 7 dpi cells, infection with Spo^-^ bacteria lead to ∼50% larvae survival at 24 hpi, compared to ∼70% for the Spo^+^ subpopulation. At the same time point, larvae infected with bacteria from 7 dLB cultures exhibited ∼70% survival for Spo^-^ cells and ∼65% for Spo^+^ cells, a difference that was not statistically significant. By 96 hpi, survival was <10% for larvae infected with expo cells, ∼40% and ∼50% for Spo^-^ and Spo^+^ 7 dpi cells, respectively. Larvae infected with 7 dLB cells maintained a similar ∼5% difference between Spo^+^ and Spo^-^ subpopulations, which remained statistically non-significant (Fig. 3a). These results indicate that Spo^-^ cells recovered from insect cadavers are more virulent than Spo^+^ insect cadavers-extracted bacteria and both Spo^+^ and Spo^-^ 7 dLB cells.

**Figure 3.**
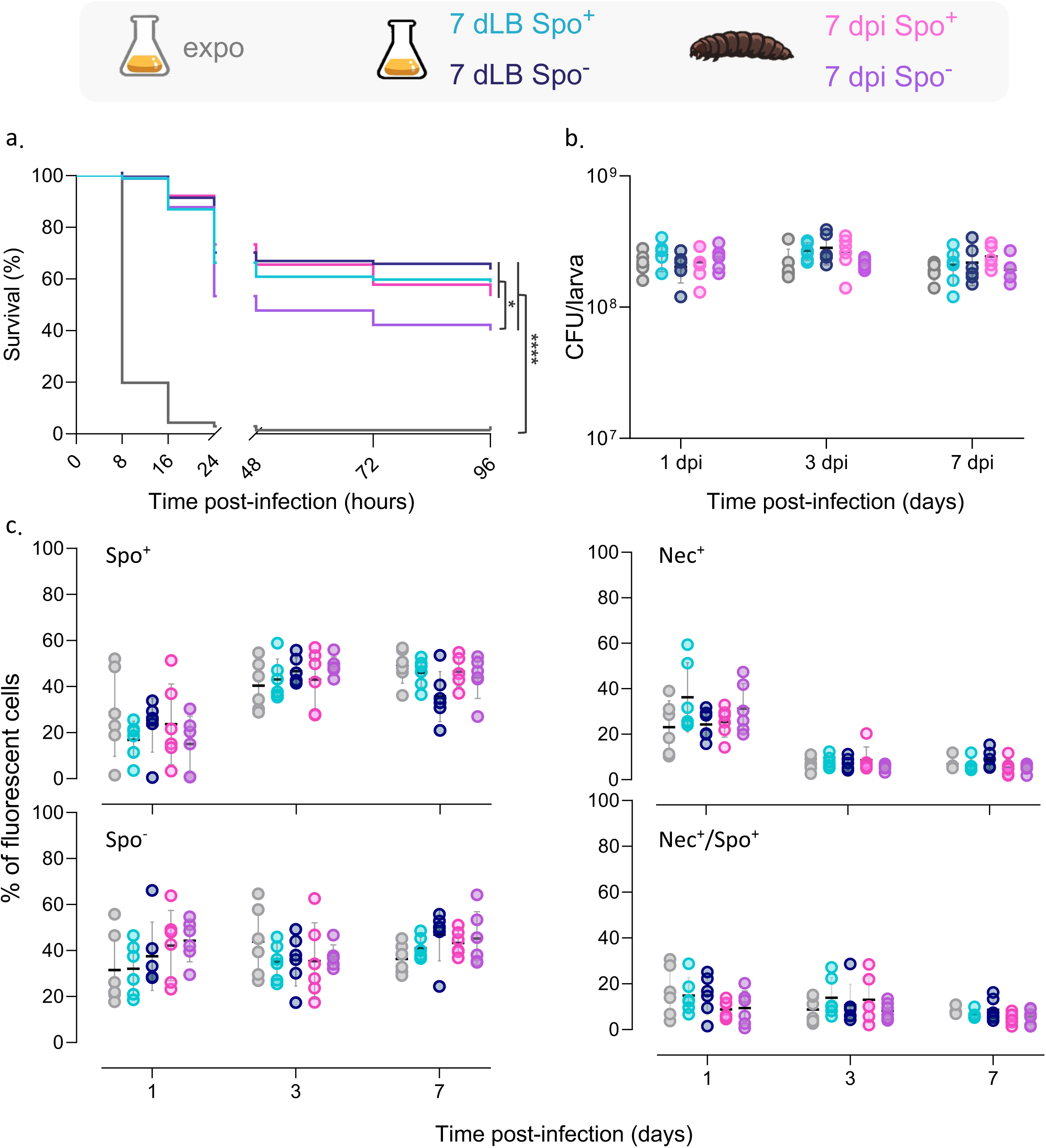
Analysis of virulence, persistence, necrotrophism and sporulation during infection with sorted bacterial subpopulations. (a) Survival of *G. mellonella* larvae following infection with sorted bacterial subpopulations from *in vitro* and *in vivo* experimental conditions using strain Bt (pP*nprA*’*gfp_Bte_AAV*-P*spoIIQ*’*mcherry*). Exponential phase cells sorted as a whole population (gray); 7 dLB cultures cells, Spo^+^ (7 dLB Spo^+^, light blue) and Spo^-^ (7 dLB Spo^-^, dark blue); 7 dpi cells, Spo^+^ (7 dpi Spo^+^, pink), and Spo^-^ (7 dpi Spo^-^, purple) were injected intrahemocoelically into third-instar larvae. 2 x 10^3^ of sorted bacteria were used for infection. Larvae were incubated at 30°C, and survival was monitored over time (hours post-infection). The data represent three independent experiments with 30 larvae per condition in each experiment. Survival curves were compared using the Gehan– Breslow–Wilcoxon test. Significance is indicated as follows: *, p < 0.05; ****, p < 0.0001. (b) Persistence of the bacteria in insect cadavers following infection with the same sorted populations described in panel a. Larvae were crushed at the indicated time points (days post-infection) using a FastPrep homogenizer and homogenates were serially diluted in saline solution and plated onto LB agar to determine colony-forming units (CFU/mL). Each symbol represents bacteria recovered from one larva (n = 6 larvae per time point, 2 larvae were used per experiment). Three independent experiments were performed. Horizontal black bars indicate the mean, and error bars represent the standard deviation (SD). (c) Flow cytometry analysis of the subpopulations after reinfection with the same sorted subpopulations described in panel a. The infection cycle dynamics were monitored with the Bt (p*PnprA*’*gfp_Bte_AAV-*P*spoIIQ*’*mcherry* strain. The percentage of fluorescent cells was determined by flow cytometry for Spo^+^ cells (top left panel), Spo^-^ cells (left bottom panel), Nec^+^ cells top right panel) and Nec^+^/Spo^+^ cells (right bottom panel) as a function of time (days post-infection). Three independent experiments were performed. Horizontal black bars indicate the mean, and error bars represent the standard deviation (SD).

We next examined persistence in insect cadavers and the ability to complete the infection cycle. Following host death, all bacterial subpopulations, whether Spo^+^ or Spo^-^ from LB or insect cadavers, persisted at a comparable level (∼2 × 10^8^ CFU/mL) over 1, 3, and 7 days post-infection (Fig. 3b). Using the *PnprA*’*gfp_Bte_AAV* and Ps*poIIQ*’*mcherry* transcriptional reporters, we observed that necrotrophism was initiated for all the infection conditions in ∼40% of the bacteria at 1 day post-infection, followed by engagement of approximately half of the population in sporulation, irrespective of their origin or sporulation state prior to infection (Fig. 3c).

Together, these results show virulence differs depending on the infecting subpopulation. However, once the host dies, bacteria persist at a similar level and undergo the infection cycle using necrotrophism as an early survival strategy followed by sporulation for part of the population.

### Bacterial tolerance to oxidative stress is enhanced in insect-extracted populations

Previous work showed that the insect cadaver represents an oxidizing environment for bacteria [26]. To investigate whether prior exposure to the insect environment influences oxidative stress adaptation, we compared tolerance to hydrogen peroxide (H₂O₂) of expo, 7 dpi and 7 dLB cells by plating the bacteria on LB agar supplemented or not with increasing concentrations of H₂O₂ using strain Bt (pP*nprA*’*gfp_Bte_AAV*-P*spoIIQ*’*mcherry*). At 0.075 mM H₂O₂, survival of 7 dpi and 7 dLB cells was markedly higher than that of expo cells (∼70% and ∼45%, respectively, compared with less than 2% for exponential-phase cells). At 0.15 mM H₂O₂, approximately 20% of 7 dpi cells survived, whereas survival was below 2% for both expo and 7 dLB cells (Fig 4.a). These results indicate that bacteria that have persisted for seven days, particularly in the insect environment, exhibit enhanced tolerance to oxidative stress.

**Figure 4.**
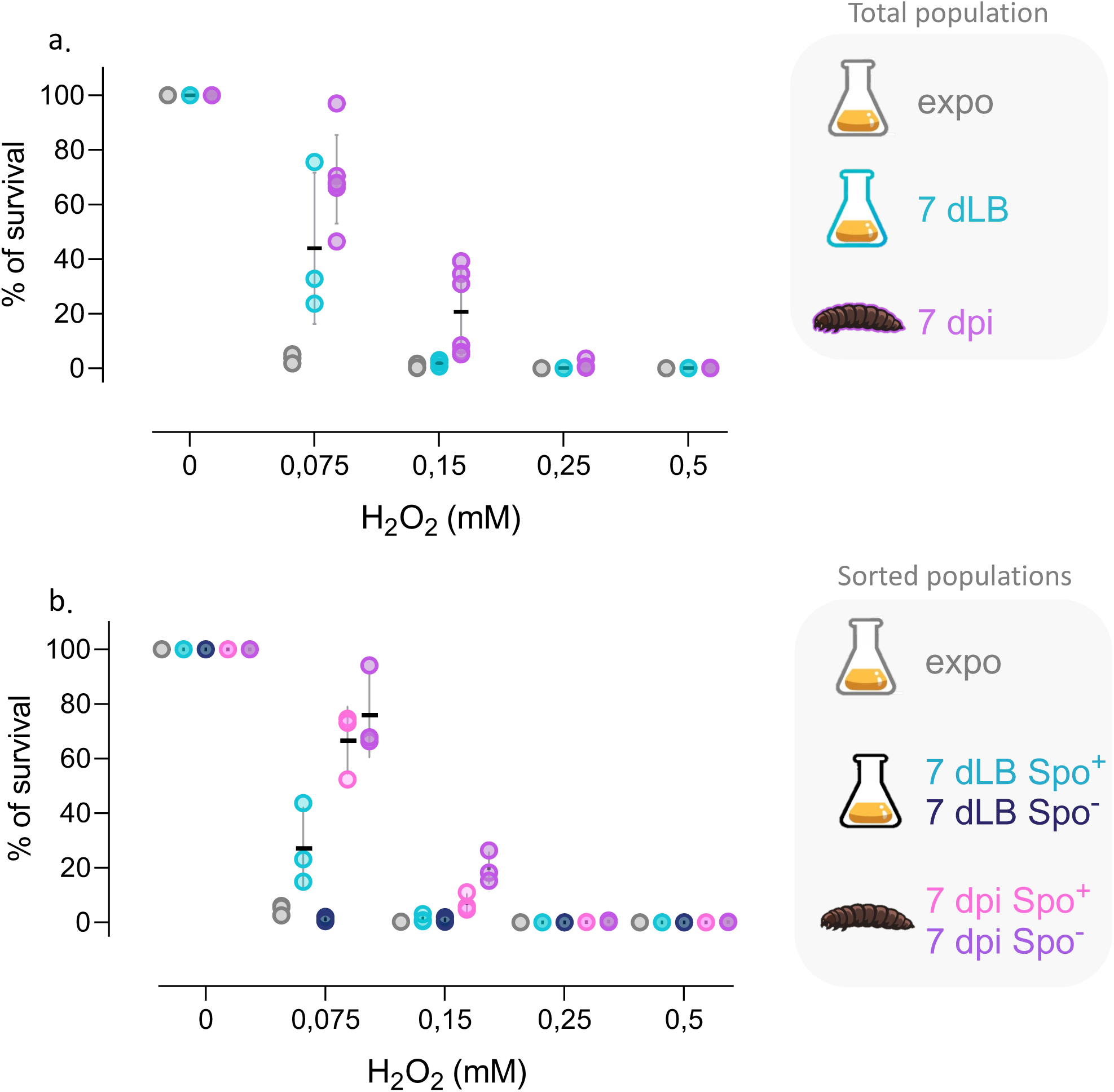
Hydrogen peroxide tolerance of total and sorted subpopulations from *in vitro* and *in vivo* conditions. (a) Hydrogen peroxide tolerance of total bacterial populations using strain Bt (pP*nprA*’*gfp_Bte_AAV*-P*spoIIQ*’*mcherry*). Bacteria from exponential phase cultures (gray), 7 dLB cultures (blue), and 7 dpi insect cadavers (purple) were serial diluted in saline and plated on LB agar supplemented or not with increasing concentrations of H₂O₂ (0.075, 0.150, 0.250, and 0.500 mM). Survival was calculated relative to untreated controls. (b) Hydrogen peroxide tolerance of sorted subpopulations using strain Bt (pP*nprA*’*gfp_Bte_AAV*-P*spoIIQ*’*mcherry*). Spo^+^ and Spo^-^ cells were sorted by FACS from 7 dLB and 7 dpi populations. Exponential phase cells were recovered as a whole population after undergoing sorting to control for potential effects of the procedure. Exponential phase (gray), 7 dLB Spo^+^ (light blue), 7 dLB Spo^-^ (dark blue), 7 dpi Spo^+^ (pink), and 7 dpi Spo^-^ (purple) cells were serial diluted in saline and plated on LB agar supplemented with the same increasing concentrations of H₂O₂, and survival was determined relative to untreated controls. Each symbol represents one biological replicate (n = 3). Horizontal black bars indicate the mean, and error bars represent the standard deviation (SD).

To examine the contribution of the different subpopulations to H₂O₂ tolerance, FACS sorting was used to isolate Spo^+^ and Spo^-^ cells from 7 dpi insect cadavers and 7 dLB cultures. Expo cells were recovered as a whole population after undergoing sorting to control for potential effects of the procedure. Exposure to 0.075 mM H₂O₂ revealed that about 25% of Spo^+^ 7 dLB cells survived, compared with 65% of Spo^+^ 7dpi and 75% of Spo^-^ 7dpi cells. At 0.15 mM H₂O₂, approximately 20% of Spo^-^ 7dpi cells and 10% of Spo^+^ 7dpi cells survived, whereas survival remained below 2% for both expo and 7 dLB populations (Fig 4.b).

Together, these data demonstrate that both Spo^+^ and Spo^-^ 7 dpi cells exhibit a higher tolerance to H₂O₂ compared to Spo^+^ or Spo^-^ 7 dLB and expo cells, highlighting the oxidative environment adaptation of the insect-extracted bacterial population.

### Non-sporulating cells outcompete spores in insect-extracted populations but are disadvantaged in 7-day LB cultures

To compare the fitness of Spo^+^ and Spo^-^ cells, competition assays were performed between subpopulations sorted by FACS from 7 dLB cultures and from 7 dpi in insect cadavers. Within each condition, Spo^+^ and Spo^-^ cells were competed against each other. Spo^-^ bacteria were isolated using the CS strain, which harbors a constitutive GFP reporter (promoter region of the *sarA* gene, P*c*’*gfp_Bte_*) and a sporulation reporter (P*spoIIQ*’*mcherry*), as GFP-positive and mCherry-negative cells. Spo^+^ bacteria were isolated as mCherry-positive cells using the S strain, which carries only the sporulation reporter P*spoIIQ*’*mcherry*. Sorted Spo^-^ (green) and Spo^+^ (red) cells were mixed at a 1:1 ratio. Mixtures were either incubated in LB for 16 h or injected into third-instar larvae and recovered at 16 h post-infection. The proportions of GFP-positive cells (derived from Spo^-^ cells) and non-fluorescent cells (derived from Spo^+^ cells that had germinated and lost the mCherry fluorescence) were then determined (Fig. 5a). As a control, expo cells from both reporter strains were passaged through the cell-sorter and mixed using the same procedure. In this condition, GFP-positive cells were competed against non-fluorescent cells, as no sporulation had yet occurred (Fig S1). The competitive index (CI) was calculated as the number of GFP-positive bacteria (Spo^-^ cells) relative to non-fluorescent bacteria (Spo^+^ cells). *In vitro*, 7 dLB Spo^-^ cells were disadvantaged compared to Spo+ cells, whereas 7 dpi Spo^-^ cells showed a slight advantage over Spo^+^ cells (Fig. 5b). *In vivo*, these trends were similar, however, there was a more pronounced advantage for 7dpi Spo^-^ cells (Fig. 5c). Pairwise comparisons using Dunn’s test revealed a significant difference between 7 dLB and 7 dpi samples (p < 0.05) (Fig.5b and c). Competitions with expo cells yielded CI values close to 1, indicating that the fluorescent constructs and the sorting procedure did not introduce a measurable fitness bias (Fig S1). These results indicate that 7 dpi Spo^-^ bacteria are better able to resume growth in a favorable condition than Spo^+^ bacteria of the same origin.

**Figure 5.**
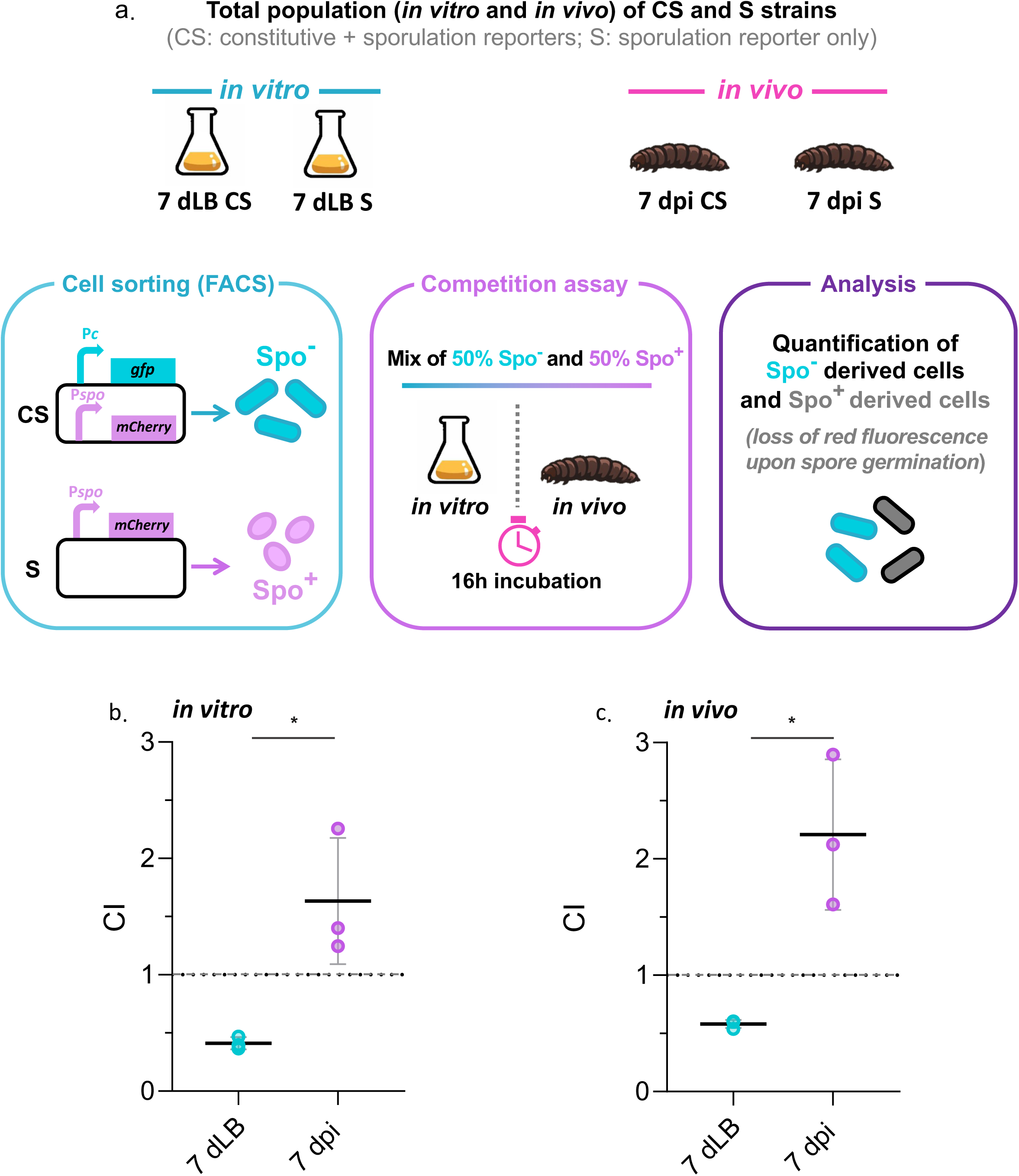
Competition assays between Spo^+^ and Spo^−^ subpopulations under *in vitro* and *in vivo* conditions. (a) Experimental design of competition assays. Spo^+^ and Spo^-^ cells were sorted by FACS from 7 dLB cultures and 7 dpi insect cadavers using Bt (pP*c*’*gfp_Bte_*-P*spoIIQ*’*mcherry*) and Bt (pP*spoIIQ*’*mcherry*) strains. Spo^-^ cells were isolated using the CS strain carrying a constitutive GFP reporter and a sporulation reporter. GFP-positive and mCherry-negative events were sorted as Spo^-^ cells. Spo^+^ cells were isolated as mCherry-positive events using the S strain carrying only the sporulation reporter. Sorted Spo^-^ (green) and Spo^+^ (red) cells were mixed at a 1:1 ratio. Mixtures were either inoculated in LB and incubated for 16 h (*in vitro*) or injected into third-instar *G. mellonella* larvae and recovered 16 h post-infection (*in vivo*). The proportions of GFP-positive cells (derived from Spo^-^ cells) and non-fluorescent cells (derived from Spo^+^ cells that had germinated and lost mCherry fluorescence) were determined, and the competitive index (CI) was calculated as described in Material and Methods. (b) *In vitro* competition assays. Sorted samples derived from 7 dLB cultures, and 7 dpi insect populations were mixed (1:1 ratio) and inoculated in LB. Spo^-^ competitive indexes were determined after 16 h of incubation. (c) *In vivo* competition assays. Sorted samples from 7 dLB cultures, and 7 dpi insect populations were mixed (1:1 ratio) and injected into third-instar *G. mellonella* larvae and recovered at 16 h post-infection. Spo^-^ competitive indexes were determined at 16 h post-infection. Each symbol represents one biological replicate (n = 3). Horizontal black bars indicate the mean. Statistical analysis was performed including expo cells using a Kruskal–Wallis test followed by Dunn’s multiple-comparison test. Significance is indicated as follows: *, p < 0.05.

### Insect-extracted spores germinate more efficiently and present lower heat resistance than *in vitro*-grown spores

To further investigate how spore formation in the insect environment affects their properties, we measured heat resistance and germination dynamics of spores produced under different conditions from strain Bt (pP*spoIIQ*’*mcherry*). Spores extracted from 7 dpi insect cadavers, 7 dLB cultures, and 7 day HCT sporulation medium (7 dHCT) were exposed to 90°C for varying durations. Surviving spores were quantified as colony-forming units per mL (CFU/mL) on LB agar, and survival curves were expressed as log(N/N₀), where N₀ is the initial spore concentration and N the number of surviving spores after heat treatment. Figure 6a shows that 7 dHCT spores were the most heat-resistant (log(N/N₀) ≈ −2.5 at 30 min), 7 dLB spores had lower resistance (≈ −4), and 7 dpi spores were the least heat-resistant (≈ −4.5). Survival remained stable at 60 min. These results indicate that the environmental conditions during sporulation strongly influence spore thermal tolerance and that the insect cadaver environment induces the formation of spores that are less heat-resistant than 7 dHCT and 7 dLB spores.

**Figure 6.**
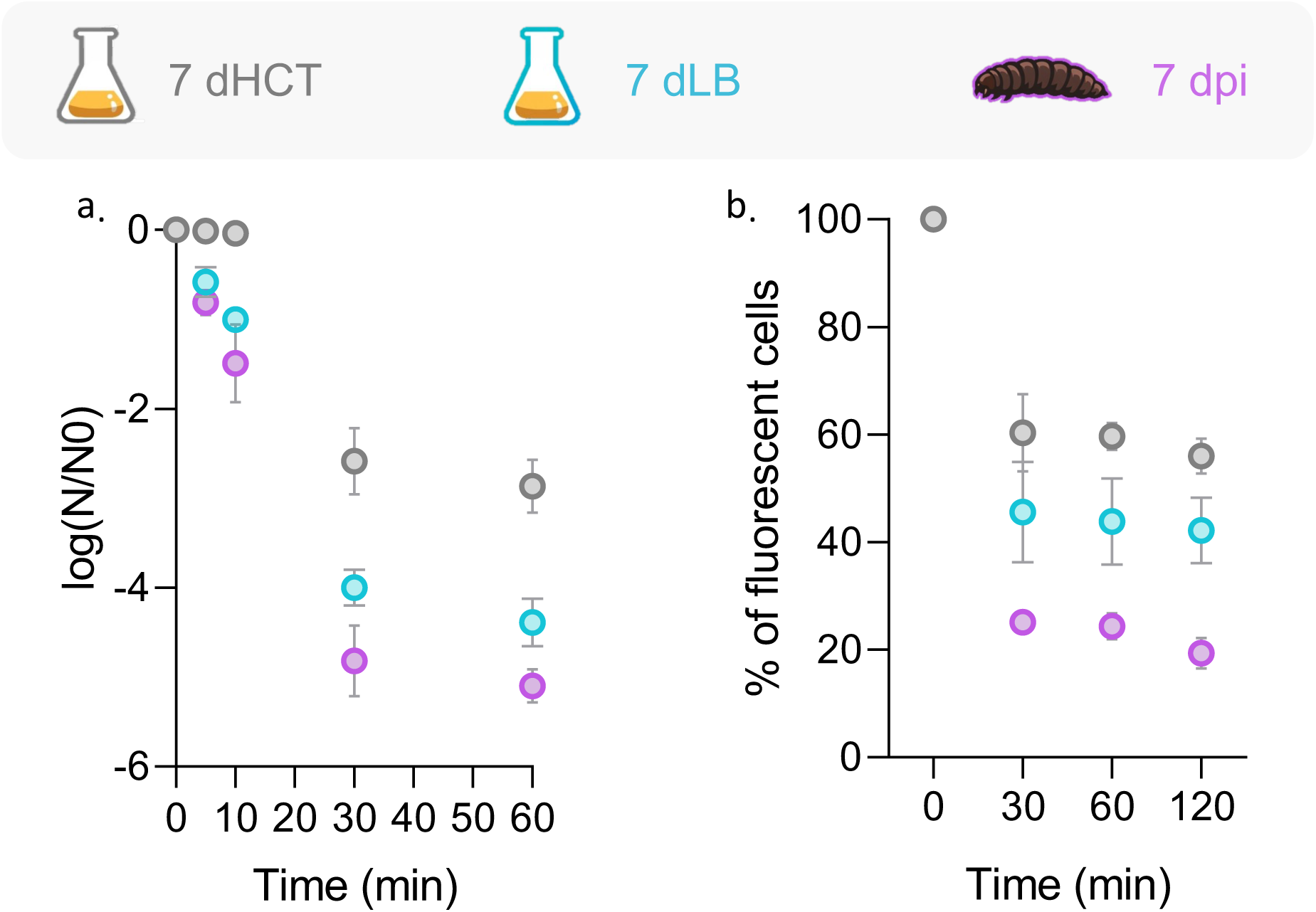
Thermal resistance and germination kinetics of spores produced under *in vitro* and *in vivo* conditions. (a) Heat resistance of spores produced under *in vitro* and *in vivo* conditions. Using Bt (pP*spoIIQ*’*mcherry*) strain, spores prepared from 7 dHCT cultures (gray), 7 dLB cultures (blue), and 7 dpi insect cadavers (purple) were exposed to 90°C for the indicated times. Heat-resistant spores were quantified by plating onto LB agar and the data expressed relative to the initial spore concentration, allowing comparison of survival kinetics over time. Data are presented as the mean of three independent experiments, and error bars represent the standard deviation (SD). (b) Germination dynamics monitored using the P*spoIIQ*’*mcherry* reporter. Spores prepared from 7 dHCT (gray), 7 dLB (blue), and 7 dpi (purple) conditions were incubated in germination buffer supplemented with L-alanine, inosine, and ten fold diluted LB + 3% glucose as described in Material and Methods. The proportion of Spo^+^ was quantified by flow cytometry at the indicated time points and normalized to the initial value (t = 0) to compare germination kinetics between conditions. Data represent the mean of three independent experiments, and error bars indicate the standard deviation (SD).

Then, germination dynamics were measured using the Bt (pP*spoIIQ*’*mcherry*) strain. Indeed, spores present a red fluorescence that is lost at the start of germination when cells start to elongate. Spores were incubated in Tris-NaCl-based germination buffer supplemented with the L-alanine and inosin germinants and ten-fold diluted LB with 3% glucose. In this condition, diluted LB+3% glucose permits cell elongation without division, ensuring that the loss of the mCherry signal reflects germination of spores rather than proliferation of new cells. The proportion of fluorescent cells (Spo^+^) was quantified at 30, 60, and 120 min, normalized to t = 0 (Fig. 6b). Most germination occurred within the first 30 min, with ∼25% of remaining Spo^+^ cells for 7 dpi spores, ∼45% for 7 dLB spores, and ∼60% for 7 dHCT spores. These data show that spores produced in the host exit dormancy more efficiently than *in vitro*-produced spores. Together, these results demonstrate that host-extracted spores exhibit better germination and lower heat resistance.

## Discussion

Our study shows that the *B. thuringiensis* populations originating from insect cadavers exhibit distinct physiological traits such as a higher proportion of non-sporulating cells than *in vitro* grown bacteria, and also present a reduced cell length compared to the latter. This result indicates that this environmental context influences cell morphology in addition to differentiation. Previous work on *Bacillus* has highlighted that heterogeneous populations coexists in complex environments such as biofilms and insect host, suggesting that the environment can shape population structure and subpopulation dynamics [25, 32, 33, 26]. The morphological differences observed here may reflect adaptive shifts in cellular physiology under host-derived stress conditions. In other bacterial species, oligotrophic or stress-induced states are associated with reduced cell size and altered metabolic activity, consistent with survival strategies under nutrient limitation or environmental challenge [34, 35, 14, 36].

Despite being less virulent than exponential-phase cells, bacteria recovered from insect cadavers were more virulent in the *G. mellonella* infection model than those cultivated for 7 days in LB. This suggests that prior host exposure primes bacteria for enhanced virulence. Such priming is consistent with reports that bacteria can establish an induced memory response during infection, which increases colonization efficiency and pathogenicity [37; 38]. Exponential-phase cells are highly virulent likely because of immediate metabolic activity and rapid division leading to the production of virulence factors. In contrast, 7 dLB or 7 dpi cells show a brief initial growth delay compared to expo cells, visible as smaller colonies that reach final similar size to those of exponential-phase cells [26]. Interestingly, once the host was dead, all bacterial populations, regardless of their origin, persisted at a similar level and followed the canonical infection cycle of necrotrophism and sporulation. Notably, non-sporulating cells gave rise to the same subpopulations as exponential-phase bacteria, indicating they are capable of sporulation and do not result from a sporulation-null mutation. The existence of this subpopulation may either reflect phenotypic plasticity linked to environmental constraints or a bet-hedging strategy. The competition assays showed that Spo^-^ cells outcompeted Spo^+^ cells for growth when both extracted from insects, whereas 7 dLB Spo^-^ cells did not outcompete 7 dLB Spo^+^. Such competitive differences align with documented strategies in sporulating bacteria, where phenotypic heterogeneity within an isogenic population provides fitness advantages in fluctuating environments. For instance, delayed commitment to sporulation in *B. subtilis* allows a subpopulation to continue vegetative growth during environmental uncertainty, a form of adaptive bet-hedging [39]. Sporulation itself represents a major energetic investment, as the production of a spore requires extensive resource allocation and divertes metabolites from growth [40]. Similarly, *B. subtilis* committing to sporulation can carry an immediate fitness cost if nutrient limitation is transient, as non-sporulating cells are better positioned to exploit sudden nutrient availability [41]. In contrast, 7 dLB Spo^-^ cultures do not outperform co-occurring spores. Extended stationary phase in nutrient-depleted LB induces physiological changes and intracellular damage that delay regrowth, producing heterogeneous recovery [42]. Although these 7 dLB Spo^-^ cells remain viable, they show slightly lower virulence and reduced resistance to hydrogen peroxide compared with 7 dLB Spo^+^ cells, possibly limiting their competitive advantage. 7 dpi (Spo^+^ and Spo^-^) cells, however, exhibit higher tolerance to H₂O₂, suggesting that this environmental history prepares these bacteria for host adaption. This pattern is consistent with prior work showing that spore resistance to chemical insults depends on sporulation conditions. In *Bacillus atrophaeus*, resistance to H₂O₂ varies with sporulation parameters such as pH and temperature, underscoring the role of sporulation context in determining spore robustness [43]. In *G. mellonella*, reactive oxygen species including hydrogen peroxide are produced in the hemolymph during infection [44, 26], and previous data show oxidative stress response genes expression in bacteria extracted from cadavers compared with LB-grown cells in *B. thuringiensis* [26], indicating that Spo^-^ cells encounter and respond to an oxidative environment *in vivo*.

In our study, spores formed in a rich sporulation medium (HCT) exhibited the highest wet-heat resistance (90 °C) and germinated more slowly under permissive conditions, whereas insect-derived spores, were less heat-resistant and initiated dormancy exit more efficiently. This trade-off between stress resistance and germination mirrors previous findings that sporulation conditions shape both spore composition and behavior. Structural and compositional features conferring high thermal resistance, while enhancing survival, can inherently slow or reduce dormancy exit. For instance, medium composition and sporulation environment modulate germination kinetics and resistance in *Bacillus spp.* [45]. Highly heat-resistant *Bacillus thermoamylovorans* spores also germinate poorly despite functional germinant receptors [46]. Moreover, sporulation timing generates phenotypic heterogeneity in revival capacity as spores formed earlier during starvation germinate more efficiently than later-forming spores [47; 48]. The comparison of *in vitro-* and insect-derived spores highlights differences in heat resistance and germination dynamics.

Comparison with previous studies highlights the importance of the ecological context. While our data indicate that a significant fraction of host-derived spores initiate germination quickly under permissive conditions, other work showed that spores recovered from 10-20-days-old cadavers of another insect species can display reduced germination compared with laboratory spores, implying that pro-longed host exposure may decrease germination in some contexts [49]. Such discrepancies may arise from differences in the timing of spore extraction or cadaver age and insect species, underscoring that the host environment has complex and time-dependent effects on spore physiology.

Collectively, our findings show that passage through a natural insect host reshapes the population structure of *Bacillus thuringiensis*, promoting the persistence of distinct sporulating and non-sporulating subpopulations with contrasting virulence, stress tolerance and competitive abilities. Spo^-^ cells from insect cadavers outcompete Spo^+^ cells and show higher virulence, while host-derived spores germinate faster than in vitro-grown spores, suggesting that these traits confer fitness advantages for persistence in fluctuating environments and better growth when conditions improve. These results support a framework in which the ecological context of sporulation and host association shape the functional traits of *Bacillus* subpopulations. Future work should elucidate the molecular bases governing entry into and exit from distinct survival programs, as well as stress responses, in host-derived bacteria.

### Data availability

The data underlying this article are available in the article and in its online supplementary material.

## Acknowledgements

This work was supported by the ANR (French National Research Agency) funding PHHASt (ANR22-CE20-0013) and by the MICA department of INRAE (French National Research Institute for Agriculture, Food and Environment).

The present work has benefited from the Imagerie-Gif core facility supported by the ANR (FBI ANR-24-INBS-0005 (BIOGEN); SPS ANR-17-EUR-0007, EUR SPS-GSR) and with financial support from ITMO Cancer of Aviesan and INCa on funds administered by Inserm. It has also benefited from the IMI (Imagerie et Infectiologie) facility, INRAe Val de Loire. We are grateful to Stéphane Perchat and Emilie Verplaetse for helpful discussions.

## Author information

### Contributions

Conceptualization: H.T. and L.S. ; Methodology: H.T. and L.S.; Investigation: H.T. Insect reinfection : H.T. and C.B. Cell Sorting : Y.L., A.S., M.B. ; Visualization: H.T.; Funding acquisition: L.S.; Project administration: L.S.; Supervision: L.S.; Writing – original draft: H.T.; Writing – review and editing: H.T. and L.S..

