## supplemental material for "Environmental history shapes host-associated dynamics of sporulating and non-sporulating bacterial subpopulations during infection"

The authors declare that there are no competing interests in relation to the work described.

**MATERIALS AND METHODS**

**DNA manipulations**

Plasmid DNA was extracted from *E. coli* by a standard alkaline lysis procedure, using a Promega kit (Promega, Madison, Wisconsin USA). Restriction enzymes, T4 DNA ligase, Standard Taq DNA polymerase and Phusion high-fidelity DNA polymerase were purchased from New England Biolabs (Ipswich, MA, USA) and used as recommended by the manufacturer. The oligonucleotide primers (Table S1.c) used for PCR amplification were synthesized by Eurofins Genomics (Nantes, France). PCR was performed with a 2720 Thermak cycler (Applied Biosystems). All constructs were systematically verified by PCR followed by sequencing of the region of interest. Nucleotide sequences were determined by Eurofins Genomics (Köln, Germany).

**Flow cytometric analysis**

For GFP-based fluorescence, a solid blue-laser emitting at 488 nm was used, combined to a 500-nm long pass dichroic mirror and a 527-nm band pass filter (512–542) (FL1 Channel). For mCherry-based fluorescence, a solid yellow-laser emitting at 561 nm was used, combined to a 610-nm long-pass filter (FL4 channel). The analyses were performed using logarithmic gains and detector settings, adjusted on a sample of reporterless cells, to define cellular autofluorescence. Gating on FSC⁄SSC was used to discriminate bacteria from the background. For each sample, 20000 gated events were measured. Data were collected with the FlowMax software (Sysmex Partec, France) and analyzed with the Weasel 3.3.3 software (WEHI, USA).

To identify positive and negative populations on FL1/FL4 bi-parametric cytograms, we applied the 98% division line for each fluorescent marker, i.e. we set the threshold on the reporterless strain so that 98% of the population gave a fluorescence intensity below the threshold. Bacteria with a fluorescence signal above the threshold were considered positive. We cannot exclude that for a reporter expressed at a low level, a few positive cells might have a fluorescent intensity similar to that of the reporterless cells and be included in the negative population.

**Figure S1**

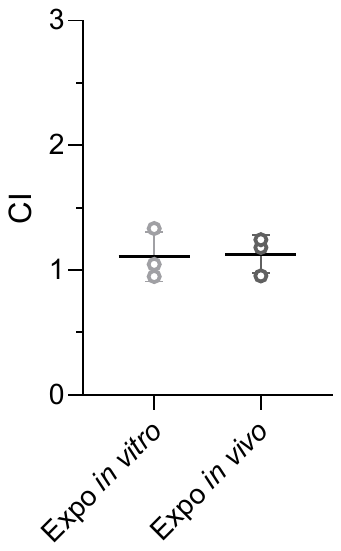

**Fig S1. Competition assays between CS and S strains under *in vitro* and *in vivo* conditions.** Using CS strain Bt (pP*c*’*gfp*_Bte_-P*spoIIQ*’*mcherry)* and S strain Bt (pP*spoIIQ*’*mcherry),* expo cells grown in LB at 30°C were mock-sorted using FACS. Cells from each strain were mixed (1:1 ratio) and inoculated in LB (*in vitro)* or injected into third-instar *G. mellonella* larvae (*in vivo)*. Competitive indexes were determined after 16 h incubation in LB or at 16 h post-infection of *G. mellonella*. Each symbol represents one biological replicate (n = 3). Horizontal black bars indicate the mean.

**Table S1.a. Plasmids used in this study.**

| **Name** | **Relevant information** | **Référence** |
| --- | --- | --- |
| pHT304 | Replicative multicopy *E. coli*/*B. thuringiensis* shuttle vector. | (1) |
| pHT304.18 | Replicative multicopy *E. coli*/*B. thuringiensis* shuttle vector. | (2) |
| pP*spoIIQ*’*mcherry* | pHT304.18 vector harboring a transcriptional fusion between *B. thuringiensis* codon optimized *comGAmcherry* and the *spoIIQ* promoter region to assess sporulation. | (3) |
| pP*nprA*’*gfp_Bte_AAV-*P*spoIIQ*’*mcherry* | pHT304.18 vector harboring transcriptional fusions between the *nprA* promoter region and the *B. thuringiensis*-codon optimized *gfp_Bte_AAV* (encoding an unstable Gfp) and the *spoIIQ* promoter region and the *B. thuringiensis*-codon optimized *comGAmcherry*, to monitor necrotrophism and sporulation | (3) |
| p*PspoIIQ*’*mcherry*-P*c*’*gfp_Bte_* | Pc (the promoter region of the *sarA* gene) was amplified by PCR from plasmid pHT304-18-P*sarA*’*gfp_Bte_* (laboratory stock, unpublished) using primer pairs P*sarA*-F-SalI/P*sarA*-R-AscI and cloned between the SalI and AscI restriction sites of pP*spoIIQ*’*mCherry*-*gfp_Bte_AAV* to obtain pP*spoIIQ*’*mCherry*-P*sarA*’*gfp_Bte_AAV*. The *gfpAAV* sequence (encoding an unstable Gfp) was then digested with AscI and EcoRI and replaced with the *B. thuringiensis*-codon optimized *gfp_Bte_* (encoding a stable Gfp) digested from plasmid p*mcherry*-*gfp_Bte_* (unpublished). This construction was used to sort green non-sporulating cells and red sporulating cells. | This study |

**Table S1.b. Strains used in this study.**

| **Name** | **Relevant information** | **Référence** |
| --- | --- | --- |
| Bt (pHT304) | *B. thuringiensis* 407^-^ carrying the empty pHT304 vector and used as a negative fluorescence control. | (4) |
| Bt (pP*spoIIQ*’*mcherry*-P*nprA*’*gfp_Bte_AAV*) | *B. thuringiensis* strain 407^-^ in which we measure the activity of the promoter of *nprA*, using a reporter gene encoding an unstable *B. thuringiensis* optimized codon GFP, as well as the activity of the promoter of *spoIIQ*, using mCherry. | (3) |
| Bt (pP*spoIIQ*’*mcherry*) or S | *B. thuringiensis* strain 407^-^ in which we measure the activity of the promoter of *spoIIQ*, using a reporter gene encoding mCherry. | (3) |
| Bt (p*PspoIIQ*’*mcherry*-P*c*’*gfp_Bte_*) or CS | *B. thuringiensis* strain 407^-^ in which *gfp_Bte_* is transcribed under the control of the constitutive promoter P*c* (promoter region of the *sarA* gene) and mcherry is transcribed under the control of the promoter of *spoIIQ* to assess sporulation activity. This strain is used to sort green non-sporulating cells and red sporulating cells. | This study |

**Table S1.c. Oligonucleotides used in this study.**

| **Name** | **Sequence** |
| --- | --- |
| PsarA-F-SalI | acgcgtcgacCTGATATTTTTGACTAAACCAAAT |
| PsarA-R-AscI | tggcgcgccAAGGTACCCGGGGATCCGATGCATC |

Purple letters indicate enzymatic restriction sites.
